# Recurrent Inhibitory Dynamics in the Entorhinal Cortex Support Pattern Separation

**DOI:** 10.1101/2024.11.14.623535

**Authors:** Yicong Zheng, James Antony, Charan Ranganath, Randall C. O’Reilly

## Abstract

The entorhinal cortex (EC) provides the major input to the hippocampus (HPC). Numerous computational models on the EC propose that its grid cells serve as a spatial metric, supporting path integration and efficient generalization. However, little is known about how these cells could contribute to episodic memory, which emphasizes episode-specific representations that align with pattern separation. Taking into consideration anatomical specifications of EC inputs to the HPC and computational principles underlying the EC-HPC memory system, we argue that EC layer IIa (EC2a) supports pattern separation and EC layer IIb/III (EC2b/3) supports generalization. Utilizing recurrent inhibition and the nature of single EC2a neurons binding converging inputs from the neocortex (i.e., conjunctive coding), we built a biologically-based neural network model of the EC-HPC system for episodic memory. By examining how EC2a transformed its cortical inputs and output them to the trisynaptic pathway (EC2a - Dentate Gyrus - CA3 - CA1), we found that instead of systematically generalizing across similar inputs, recurrent inhibition and conjunctive coding in EC2a neurons support strong pattern separation and increase mnemonic discrimination. Furthermore, lesioning EC2a neurons in the model resembled memory impairments found in people with Alzheimer’s Disease, suggesting an intertwined relationship between memory and the majority of pure grid cells in the EC. On the other hand, the topographically organized monosynaptic pathway (EC2b/3 - CA1) is computationally more suitable for efficient factorization and generalization. This model provides novel anatomically-based predictions regarding the computational roles of EC cells in pattern separation and generalization, which together form a critical computational framework for both episodic memory and spatial navigation.

## Introduction

The hippocampus (HPC) is widely acknowledged to play a critical role in episodic memory and spatial navigation, based on converging evidence from studies in humans, monkeys, and rodents. Traditionally, the entorhinal cortex (EC) has been viewed as a “funnel” to the HPC, meaning it accumulates information from across the neocortex and passes it onto the HPC without much functionally-specific processing (Lavenex & Amaral, 2000; Manns & Eichenbaum, 2006; Mishkin et al., 1997; Ranganath & Ritchey, 2012; Squire & Zola-Morgan, 1991). With the discovery of grid cells in the entorhinal cortex (EC), it has become apparent that the EC also contributes to navigation, and that it is an integral component of the hippocampal system.

Although researchers have proposed that grid cells in the EC might also contribute to episodic memory (M.-B. Moser et al., 2015), their exact role remains unclear. In this paper, we present a comprehensive computational model of EC function based on detailed anatomical and physiological data, which shows how it makes significant contributions to episodic memory based on mechanisms that also give rise to grid cells, providing a unified account of EC function. We show how this model explains recent data revealing that damage to layer 2 of lateral EC (LEC), which is the first site of pathology in Alzheimer’s Disease (AD), can have significant impacts on episodic memory function.

The first clue for functions of the EC comes from its anatomical connections with the HPC. Classic “funnel” models (Lavenex & Amaral, 2000; Manns & Eichenbaum, 2006; Mishkin et al., 1997; Ranganath & Ritchey, 2012; Squire & Zola-Morgan, 1991) tend to treat the EC as a relay station that is bidirectionally connected with the HPC. However, anatomical studies of the EC-HPC projections (Nilssen et al., 2019; Steward & Scoville, 1976; Witter, Naber, et al., 2000) have long documented distinct EC inputs to the trisynaptic pathway (TSP: EC2a - Dentate Gyrus (DG) - CA3 - CA1) and the monosynaptic pathway (MSP: EC2b/3 - CA1) (Fig 1A). This suggests a more complex functional relationship between different EC layers beyond the simple funnel model. Regarding connectivity, TSP projections are more convergent, forming sparse pattern-separated representations in the DG and CA3, while MSP projections are more topographically organized, potentially useful for factorized representation (Witter, Naber, et al., 2000). This distinction is critical to understanding how the HPC could support efficient encoding and retrieval of episodic memory, as the TSP is believed to support pattern separation and pattern completion (Marr, 1971; O’Reilly & McClelland, 1994; Yassa & Stark, 2011), while the MSP may support more systematically organized memory representation for statistical learning/generalization (Schapiro et al., 2017).

**Figure 1:**
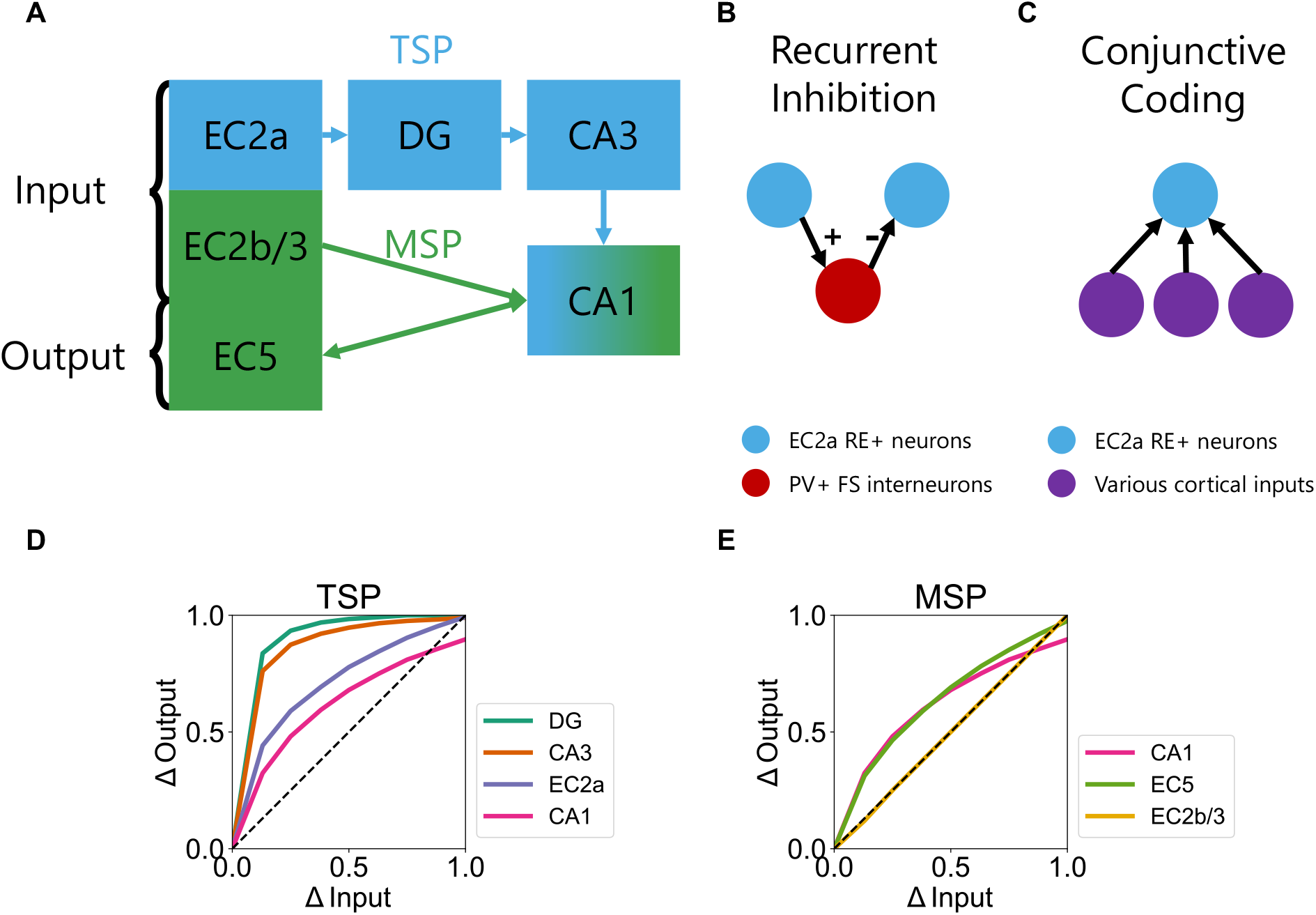
Schematic illustration of synaptic connections in the EC-HPC system. **A**. Diagram depicting the monosynaptic pathway (MSP, EC2a → DG → CA3 → CA1) and the trisynaptic pathway (TSP, EC2b/3 → CA1) and the output projection (CA1 → EC5). **B**. RE+ neurons connect with each other disynaptically via inhibitory PV+ FS interneurons, forming recurrent inhibition in the EC2a layer (Couey et al., 2013). **C**. RE+ neurons receive convergent cortical inputs from different areas, forming conjunctive coding (Canto et al., 2012; Doan et al., 2019; Nilssen et al., 2022). **D**. Pattern separation curves for the subregions of the trisynaptic pathway (TSP). Δ*Input* represents input similarities between lure and target patterns in the Input layer, while Δ*Output* indicates the similarity of neural representations in specific TSP layers for lures and targets. Lines situated in the top left half of the figure signify that output similarities are greater than input similarities (i.e., pattern separation). **E**. Pattern separation curves for the subregions of the monosynaptic pathway (MSP). EC, entorhinal cortex; HPC, hippocampus; DG, dentate gyrus; RE, Reelin; PV, Parvalbumin; FS, fast-spiking.

Another avenue to understanding EC functions comes from Continuous Attractor Network (CAN) models that shed light on how recurrent inhibition in EC2a could support the formation of grid cells during open-field foraging (Burak & Fiete, 2009; Couey et al., 2013; Fuhs & Touretzky, 2006; Guanella et al., 2007; McNaughton et al., 2006; Zutshi et al., 2018). In the anatomically based CAN model (Couey et al., 2013), recurrent lateral inhibitory connections (Fig 1B) between MEC2a Reelin+ cells create stable attractors that form grid-like relationships when receiving external inputs. While it is still unclear how this computational property of EC2a contributes to memory, its significant anatomical position as the sole EC input to the HPC signals its potential importance. Computationally, inhibition is a core ingredient for pattern separation in episodic memory models. Here, we show that strong inhibitory dynamics in the EC also drives pattern separation that improves episodic memory functions.

Although CAN models are used to explain MEC2a grid cells, the network of Reelin+ Fan cells in lateral EC (LEC) layer 2a has strikingly similar properties. Lateral EC2a is among the initial regions impacted by Alzheimer’s Disease (AD), exhibiting a significant reduction in EC2a neuron populations (Gómez-Isla et al., 1996). Reelin+ Fan cells, which are enriched in EC2a and are vulnerable to tau and beta amyloid accumulation (Igarashi, 2022; Kobro-Flatmoen et al., 2021; Stranahan & Mattson, 2010). Pattern separation and episodic memory appear to be impaired early in AD. Thus, there is reason to consider how shared computational properties of MEC and LEC can explain memory-related changes in healthy aging and AD.

To test how above mentioned features of the EC could influence its computational roles in episodic memory, we built on a biologically-based EC-HPC model for episodic memory (Zheng et al., 2022) by adding two key principles: 1) EC2a forms an inhibition-based attractor through recurrent inhibition (Fig 1B); 2) single neurons in EC2a integrate multiple cortical inputs to form conjunctive coding (Canto et al., 2012; Doan et al., 2019; Nilssen et al., 2022) (Fig 1C). Meanwhile, EC2b/3 retains a more factorized layout as the original model, given that the projections between EC2b/3 and CA1 are more topographically organized (Witter, Naber, et al., 2000). We first show that recurrent inhibition and conjunctive coding give rise to stronger pattern separation by examining model representations. Next, we show how this model can account for empirical data on decreased memory precision in aging populations, as well as AD memory deficits that have been linked to specific loss of the LEC2a Reelin+ neurons. Finally, we demonstrate that decreased EC2a recurrent inhibition in the model could lead to worse *pattern completion* capability (in addition to worse pattern separation), contradicting the prevailing thinking that the elderly have an increased pattern completion tendency (Yassa & Stark, 2011).

## Results

The RIDE (**R**ecurrent **I**nhibitory **D**ynamics in the **E**ntorhinal cortex) model is primarily developed based on our previous HPC model (Zheng et al., 2022). The architecture of the model is based on the neurobiology of the hippocampal system, where cortical inputs progress through the superficial EC, move into the TSP and MSP, and ultimately reconstruct the input patterns at the deep EC. In RIDE, we have separated the original superficial EC layer into EC2a and EC2b/3 to more accurately represent the anatomical properties of these two areas. Specifically, the original factorized structure in the superficial EC layer is retained in the new EC2b/3 layer. This layer projects to CA1 and subiculum (not modeled in the current simulation) with topographically segregated streams (Witter, Wouterlood, et al., 2000). In contrast, we added a new EC2a layer that captures the following anatomically based principles: 1) EC2a Reelin+ neurons connect to each other disynaptically via interneurons, consistent with the known recurrent inhibitory connectivity (Couey et al., 2013); 2) Single EC2a Reelin+ neurons receive convergent inputs from multiple cortical areas (Canto et al., 2012; Doan et al., 2019; Nilssen et al., 2022). We connected EC2a to DG and CA3, forming the TSP, and EC2b/3 to CA1, establishing the MSP.

### Simulation 1: EC2a Facilitates Pattern Separation as the Initial Stage of the Trisynaptic Pathway

Our initial simulation explored the mnemonic discrimination task, which requires participants to discriminate between studied (“target”) items and unstudied (“lure”) items (Kim & Yassa, 2013; Yassa, Mattfeld, et al., 2011) that vary in similarity to the targets. Discrimination between targets and highly similar lures is often hypothesized to depend heavily on hippocampal pattern separation (Stevenson et al., 2020). In the following, we simulate this task with the RIDE model and demonstrate how the anatomical structure of the EC affects pattern separation within the hippocampal system, allowing precise differentiation between lures and targets. Our model adheres to a conventional mnemonic discrimination task protocol: it initially encodes random target patterns as sequences of 0s and 1s, then computes output patterns based on these trained targets and untrained lure patterns, which are created by altering bits in the target patterns. We define target reconstruction (similar to hit) as the number of bit difference between model output and target pattern given target as input. In addition, we define lure-target mismatch (similar to correct rejection) as the number of bit difference between model output and target pattern given lure as input.

Similar to prior analyses on hippocampal functions in computational models and experiments (Guzowski et al., 2004; O’Reilly & McClelland, 1994; Wigström, 1973; Yassa, Mattfeld, et al., 2011; Yassa & Stark, 2011), we plotted the relationship between lure-target similarity (i.e., Δ*Input*) and representational similarity between lures and targets (i.e., Δ*Output*) within each hippocampal subregion (Fig 1D, E). The curvilinearity of these plots indicates the degree to which hippocampal subregions exaggerate representational differences between lures and targets (i.e., pattern separation). ^1^ EC2b/3 faithfully represents the structure of inputs, such that representational dissimilarity increases linearly with lure target dissimilarity. However, as shown in Fig 1D, differences between lures and targets are exaggerated in EC2a, as well as in DG, and CA3, consistent with previous models (O’Reilly & McClelland, 1994; Yassa & Stark, 2011). In contrast, pattern separation appears less pronounced in CA1, likely due to the direct input from EC2b/3 (Fig 1E).

As noted above, the key differences between the current model and previous computational models of the HPC are recurrent inhibition and conjunctive coding (or, convergence of inputs) in EC2 neurons. Thus, our next analyses characterized the computational significance of these two factors on pattern separation both within EC and within different hippocampal subregions.

We systematically manipulated the strength of recurrent inhibition between cells in EC2a and examined its impact on pattern separation in the entire TSP. The theory is that in EC2a, pattern separation may arise from a sparser neural representation when recurrent inhibition is high, since using fewer neurons for memory encoding might minimize overlap between different memories. Results suggest that increasing recurrent inhibition enhanced pattern separation in the whole TSP (Fig 2A). To further examine pattern separation quantitatively, we calculated the lure-target mismatch index as defined before. This measure essentially tells how good the model is at differentiating similar patterns from the original pattern as an end result. Results showed that increasing EC2a recurrent inhibition increases lure-target mismatch (Fig 2B), indicating higher and higher pattern separation between lure and target after they go through the entorhinal-hippocampal loop.

**Figure 2:**
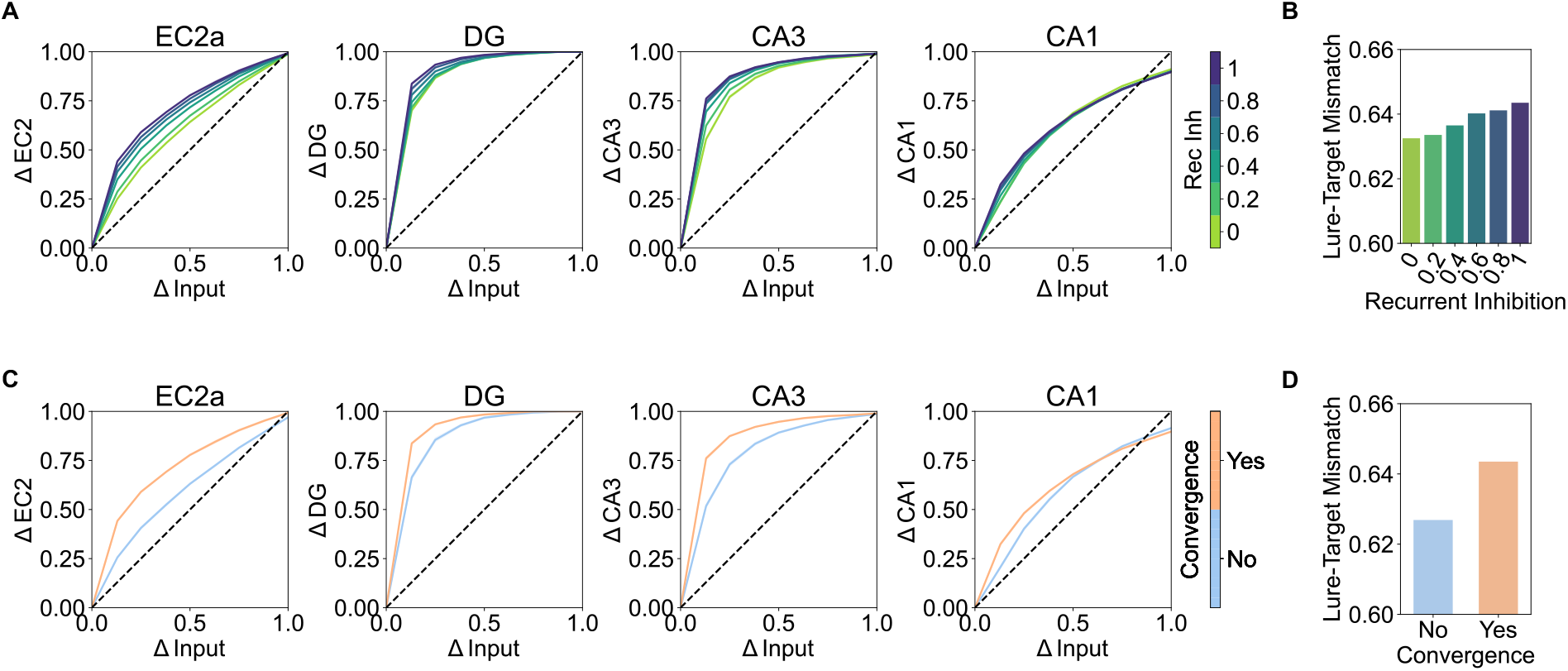
Anatomical properties of EC2a contribute to its pattern separation capabilities. **A**. Pattern separation curves across a range of EC2a recurrent inhibition strengths for the TSP. Stronger recurrent inhibition results in increased pattern separation in DG, CA3, and EC2a. **B**. Lure with higher EC2a recurrent inhibition leads to more separable EC5 output patterns from Target. **C**. Pattern separation curves for (non)convergent EC2a for the TSP. When single EC2a neurons receive convergent inputs from multiple cortical areas (i.e., multiple pools in the Input layer), the TSP exhibits enhanced pattern separation compared to when EC2a neurons do not receive convergent cortical inputs. **D**. Lure with convergent cortical inputs onto EC2a leads to more separable EC5 output patterns from Target.

Another critical biological characteristic of the EC2a is that single Reelin+ neurons in LEC2a receive converging inputs from cortical areas such as perirhinal cortex, postrhinal cortex, piriform cortex, contralateral LEC, and MEC (Doan et al., 2019; Nilssen et al., 2022), while MEC2a receive convergent inputs from presubiculum and parasubiculum (Canto et al., 2012). To examine whether such a connectivity profile could facilitate pattern separation through binding multiple elements of a memory into a conjunctive representation, we manipulated the number of neurons projecting to a single neuron in the EC2a layer in the model. We created a converging condition wherein a single EC2a neuron received inputs from multiple neurons in the Input layer, as well as a non-converging condition where a single EC2a neuron received input exclusively from one neuron in the Input layer. Results revealed that pattern separation was more pronounced with converging compared to non-converging inputs (Fig 2C). To follow up on the effect of converging vs. non-converging inputs on pattern separation, we examined lure-target mismatch to quantify the difference between model output pattern (given lure as input) and true target patterns. We found that lure-target mismatch was higher with converging inputs onto EC2a neurons than the non-converging condition (Fig 2D), suggesting that the binding property of Reelin+ neurons positions the EC2a to reduce pattern overlaps at an early stage even before the hippocampal processing loop.

Together, simulation 1 demonstrated the pattern separation nature of the EC2a, and how two of its neuroanatomical properties could underlie pattern separation in a recognition memory task. Furthermore, integrating information at the single-neuron level provides additional pattern separation. Collectively, these results offer evidence that EC2a, as the first site of the TSP, might already begin to pattern separate incoming information from the neocortex before the DG and CA3.

### Simulation 2: Neural mechanisms for declining memory in cognitive aging and Alzheimer’s Disease

Over the time course of memory declining in aging populations or people with AD, some of the earliest notable changes in the brain start within the EC2a (Igarashi, 2022), including beta-amyloid deposition (Price & Morris, 1999), neurofibrillary tangles of tau (Ziontz et al., 2019), and compromised grid-cell-like representation (Stangl et al., 2018). Importantly, Maurer et al. (2017) showed that hyperexcitability in LEC2a and CA3 cells contribute to decreased object recognition in aged poor performing rats, suggesting that the inhibition of EC2a cells could potentially lead to hyperactitve CA3 and its rigid patterns through the perforant path. To investigate the impact of age-related EC2a dysfunction on memory performance, we compared mnemonic discrimination performance of young adults using the standard RIDE model (recurrent inhibition = 1) with mnemonic discrimination in older adults by setting the magnitude of EC2a recurrent inhibition to zero.

We started by asking how young and old conditions differ in hit rates (correctly identifying targets as “Old”) and correct rejection (correctly identifying lures as “New”). By assessing the degree of match between reconstruction patterns in EC5 and the input patterns (see Methods for details), we simulated hits using Target Reconstruction and correct rejections using Lure-Target Mismatch, both defined in Simulation 1. We found that the old condition had slightly lower Target Reconstruction but significantly reduced Lure-Target Mismatch compared to the young condition (Fig 3A), suggesting greater similarity between reconstructed lure patterns and true target patterns (i.e., less pattern separation). This result pattern is in alignment with the empirical findings that older adults showed a similar level of hit rates but significantly decreased correct rejection rate compared to young adults (Fig 3A). Additionally, we computed the bias-free signal detection measure *d*^′^ (hit rate minus false alarm rate) as target reconstruction minus (1 - lure-target mismatch). This calculation demonstrated that the old model exhibited a lower *d*^′^ value (poorer discriminability), consistent with experimental data (Fig 3A). The computational evidence underscores a decrease in pattern separation in older adults, particularly within regions of the TSP (Fig 3C), supporting previous hypotheses (Yassa, Lacy, et al., 2011; Yassa, Mattfeld, et al., 2011). Altogether, these simulations suggest that reduced pattern separation in older adults could come from compromised recurrent inhibition between EC2a Reelin+ neurons.

**Figure 3:**
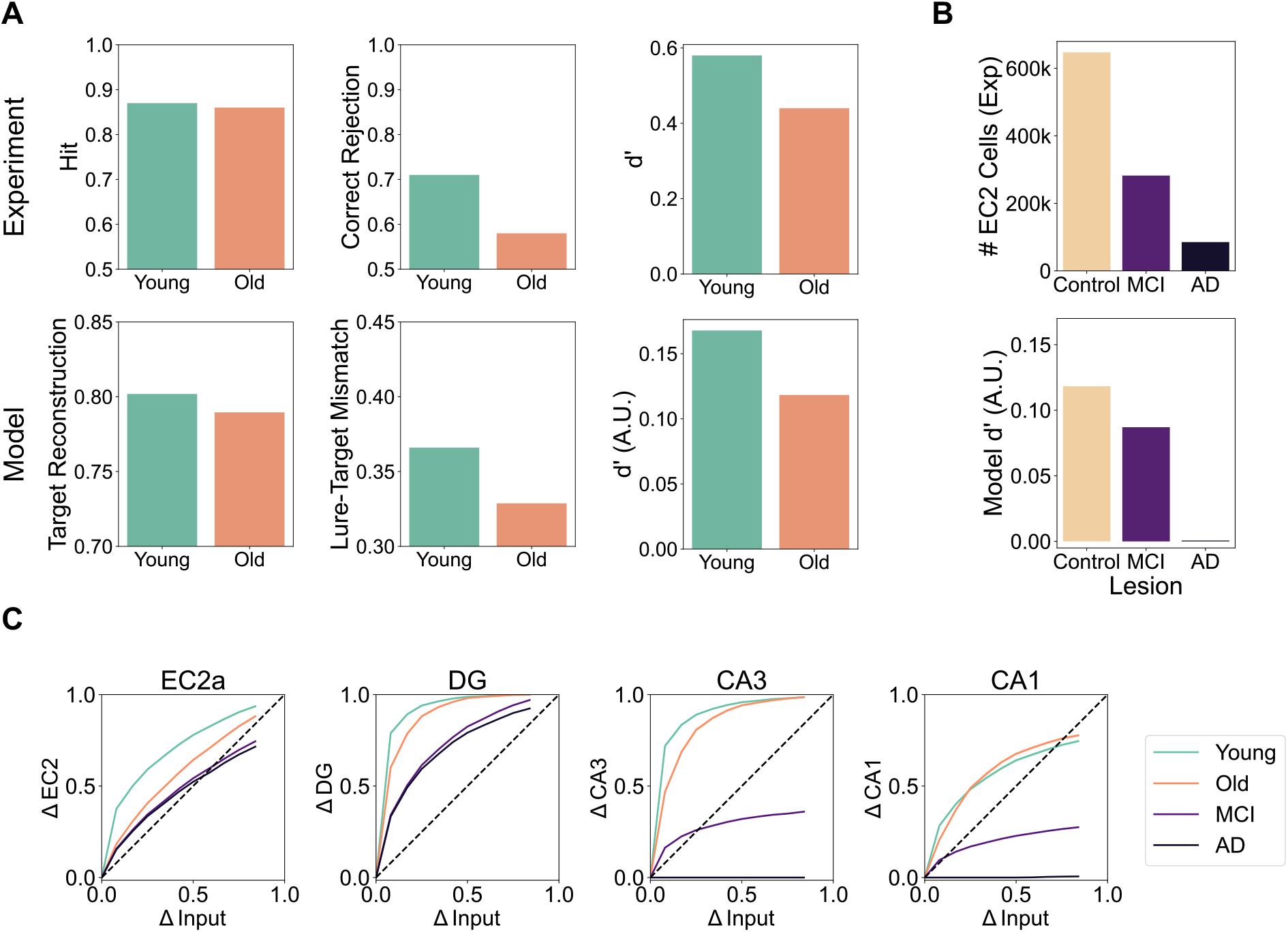
EC2a as a key to cognitive aging and Alzheimer’s Disease (AD). **A**. Experiment results for an object mnemonic discrimination task and corresponding model outcomes. Cognitively healthy older individuals exhibit slightly impaired hits and significantly impaired correct rejections. RIDE demonstrates analogous patterns of results with target reconstruction, lure-target mismatch, and *d*^*′*^ in its EC5 output layer. Note that the Young model has the same parameter as RIDE, with recurrent inhibition strength being 1, while the Old model has a recurrent inhibition strength of 0 (medium values created similar effects). **B**. EC2a neuron degeneration in Mild Cognitive Impairment (MCI) and AD populations (Gómez-Isla et al., 1996) and simulated mnemonic discrimination task performance in RIDE. EC2a neuron count decreases by 57% in MCI, and 87% in AD compared to healthy controls. When EC2a neurons are lesioned to similar levels, model performance decreases, exhibiting lower overall *d*^*′*^ values (excluding response biases). **C**. By combining the four simulated conditions in **A** and **B**, curves in the TSP show a gradual decline in pattern separation ability of the model (i.e., equivalent to cognitive decline from Young to Old to MCI to AD). Notably, in CA3, MCI fails to generate lure patterns sufficiently distinct from target patterns, suggesting a tendency to confuse targets and lures. Moreover, for AD, the CA3 and CA1 curves overlap with the x-axis, indicating that the model always generates identical patterns regardless of input similarity, which suggests a failure to retrieve any meaningful memory.

Subsequent experiments considered the potential role of EC2a Reelin+ neurons in the pathological evolution of AD. Specifically, how gradual loss of EC2a Reelin+ neurons could lead to memory changes as seen in the progression of AD. Building on the Old model with zero recurrent inhibition, we lesioned EC2a neurons (i.e., setting activity to be 0) in the model to parallel the development of AD. It is known that EC2a Reelin+ neurons are profoundly reduced in Mild Cognitive Impairment (MCI) and AD, with approximately 57% and 87% reduction of EC2a neurons compared to control in MCI and AD, respectively [Fig 3B, (Gómez-Isla et al., 1996)]. Results suggest that lesioning EC2a neurons to degrees similar to those in MCI and AD decreases *d*^′^, dramatically reducing target-lure discriminability. When EC2a neurons were nearly fully lesioned, model performance dropped nearly to zero, indicating an incapability to perform the memory task. Together, these simulations demonstrate how impaired EC2a Reelin+ neurons throughout the progression of AD could contribute to a decline in mnemonic discrimination ability.

Finally, we plotted pattern separation curves in the aforementioned simulations (Fig 3C), including Young, Old, MCI, and AD, where a similar percentage of lesions were performed on the models. Results suggest that there was a gradual decrease of pattern separation throughout the course of AD progression. Furthermore, it was in the traditional conjunctive memory hub CA3 that MCI showed a flattened pattern separation curve, meaning the lure representations in CA3 remained relatively similar to the target representations regardless of how divergent they were in the Input layer, suggesting a blur between lures and targets. Moreover, lure and target representations in AD were always almost the same as depicted by the curve that overlaped with the x-axis, suggesting that the model was not only unable to discriminate between lures and targets, but failed to relate these test cases to what it originally learned.

In summary, the simulations elucidate potential neural mechanisms driving the decline in memory with aging and AD. They accurately reflect the experimental behavioral patterns in older adults, such as the reduced rate of correct rejections. Furthermore, the simulated lesioning of EC2a neurons aligns with AD progression, particularly noting a substantial impairment in overall object discriminability when significant EC2a neuron loss occurs.

### Simulation 3: Pattern separation facilitates, but not contradicts pattern completion

Pattern separation and pattern completion have traditionally been recognized as two opposing processes (O’Reilly & McClelland, 1994). That is, given a fixed distance between two input patterns, if the output patterns’ distance is larger than from input, then a network is said to be doing pattern separation, but if the opposite is true, then the network would be said to be doing pattern completion. When applied to behavioral and neural data, the assumption has been that age-related reductions in mnemonic precision (Yassa, Lacy, et al., 2011; Yassa, Mattfeld, et al., 2011) are indicative of worse pattern separation but more pattern completion (Yassa & Stark, 2011). Our simulations suggest that this is not the case.

In computational terms, pattern completion is operationally defined as the ability for a model to recall a stored pattern from an incomplete cue, which is comparable to what is measured behaviorally in cued-recall tests. To measure pattern completion behaviorally, Vieweg et al. (2015) developed a task in which participants studied complex pictures and were then shown cue pictures that are either studied or unstudied with gradually deleted features (Fig 4A). They were instructed to respond with either “new” (indicating that they have not learned the picture) or any of the original word associated with the picture (e.g., kitchen). As shown in Fig 4B, results from the experiment showed that, relative to young adults, older adults were biased towards responding with studied words (more hit and false alarm, see Methods for details). This fits nicely with previous findings (Yassa, Lacy, et al., 2011; Yassa, Mattfeld, et al., 2011) where older adults tend to incorrectly treat lures as targets. However, when we took the literal definition of pattern completion, we found that older adults had lower hit rate, meaning they couldn’t complete the learned patterns from partial cues as accurate as young adults.

**Figure 4:**
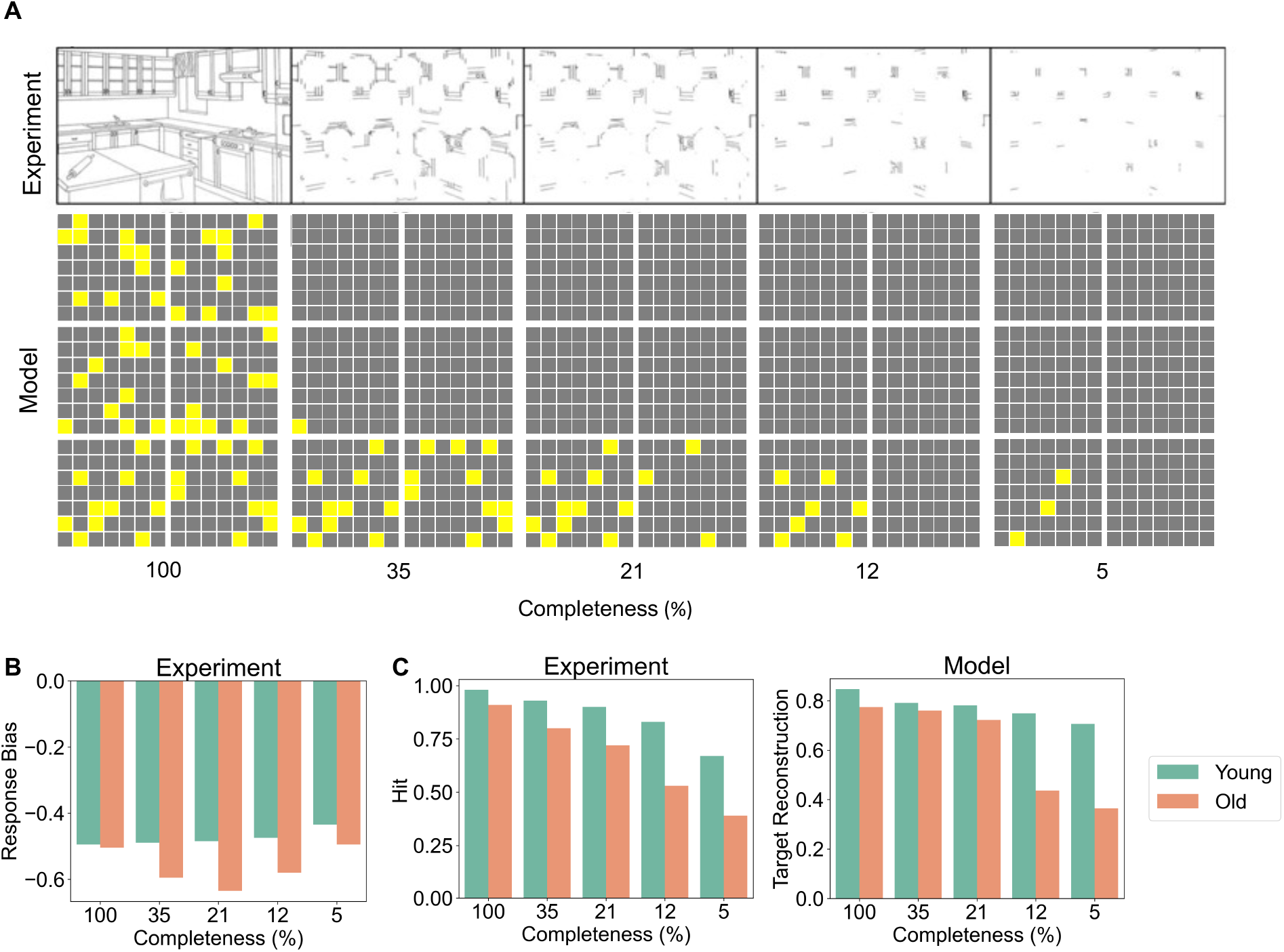
EC2a inhibition facilitates both pattern separation and pattern completion. **A**. Experiment stimuli adapted with permission from Vieweg et al. (2015) with varying degrees of completeness and corresponding modeling patterns. Subjects were instructed to respond in either one of the previously learned semantics associated with memorized figures or “new”. **B**. Reanalyzed experiment results show a response bias of −0.5 × (hit + false alarm) in older adults, indicating a tendency to respond with previously learned semantics. **C**. Hit and target reconstruction calculated from experiment data and modeling results, respectively, indicate that the capability to perform the pattern completion task declines in older adults (model) compared to young adults (model) across all completeness conditions.

We then took the Young and Old versions of the RIDE model used in Simulation 2 (with the critical difference being presence of recurrent inhibition in EC2a) and simulated the same experiment (see details in Methods). Results showed a consistent trend of decreased target reconstruction when reducing completeness in the partial cue input, and that the Old model is worse at reconstructing the target pattern given partial cue with all completeness conditions (Fig 4C), suggesting worse pattern completion capability. Considering that in the experiment older adults showed response biases towards responding with studied words, the degree of them not being able to complete the studied patterns might be more drastic, leading to even worse pattern completion capability than shown in Fig 4C. In Simulation 2, we have demonstrated that the Young model has a more pattern-separation-like curve in Fig 3C, but critically, we should not conclude that it is worse at pattern completion than the Old model. In fact, a model with better pattern separation could have better pattern completion, as shown here in Fig 4C.

Overall, this simulation provides a conclusion that contradicts common beliefs regarding pattern separation and pattern completion as mutually exclusive. Decreasing recurrent inhibition in the EC2a layer reduced pattern separation in Simulation 2, and the same manipulation also reduced the ability for the model to recall stored patterns based on partial cues. In other words, disinhibition in EC2a could be a key mechanism for decreased pattern separation and pattern completion in older adults.

## Discussion

In the current study, we found that neuroanatomical properties of the entorhinal cortex layer 2a Reelin+ neurons, which have previously been used to simulate grid cell activity in the medial EC, also have significant beneficial effects for episodic memory function, by increasing pattern separation of information that then projects into the hippocampus proper via the trisynaptic pathway. Specifically, the recurrent inhibition between EC2a Reelin+ neurons causes EC2a to form sparser patterns, resulting in more pattern separation. Furthermore, more broadly convergent (“conjunctive”) input into individual EC2 neurons produced more pattern separation and better pattern reconstruction compared to the case without convergent inputs. We were able to simulate the impairments in mnemonic discrimination as seen in older adults by decreasing recurrent inhibition. Interestingly, we found that this impairment in pattern separation also led to worse pattern completion, in contrast to some prior interpretations of data that can be reinterpreted as resulting from a change in the overall response bias (i.e., an increased bias to classify novel items as “old”) (Kim & Yassa, 2013; Stevenson et al., 2020; Yassa, Mattfeld, et al., 2011). Finally, we showed that lesioning EC2a neurons causes weaker pattern separation and impaired memory performance, as seen in the progression of AD, which has been shown to selectively target these EC2a Reelin+ neurons. Together, these results suggest an important role for EC2a in episodic memory function, based on its idiosyncratic anatomical features.

According to the complementary learning systems (CLS) framework, the hippocampus is specialized for pattern separation to support fast episodic memory encoding, whereas the neocortex uses more distributed, overlapping patterns of activity to support generalization to novel situations (McClelland et al., 1995). However, the role of the EC in this framework is unclear. At present the majority of models of EC function have focused on how grid cells could support generalization (Behrens et al., 2018; Gustafson & Daw, 2011; Momennejad, 2020), but very few models have touched on the issue of pattern separation (Kerdels & Peters, 2017). This contradiction makes it computationally interesting to examine hypotheses behind theories, how they connect, and where they differ.

The classical perspective on grid cells is that they provide a coordinate-system-like spatial metric (Bush et al., 2015; Hafting et al., 2005; E. I. Moser & Moser, 2008; Whittington et al., 2020), which depends on their ability to maintain consistent firing fields across different environments. For instance, moving from room A to a novel room B should not alter the distance between adjacent firing fields of grid cells in both rooms. However, this requirement of consistent, generalizable grid cells firing is being increasingly challenged by data showing significant differences in grid cell firing across environments (Aronov et al., 2017; Campbell et al., 2021; Low et al., 2021), which has been described as similar to the *remapping* behavior of neurons in CA3 and CA1. This remapping and context-dependence of grid cell firing is consistent with the pattern separation property of the EC2a neurons in our model. In line with our computational reasoning, Kitamura et al. (2015) found that Reelin+ cells in MEC2a form distinct representations in a novel context and drive context-specific CA3 activation, suggesting that contextual information is already encoded in EC2a Reelin+ neurons and that the entire TSP is crucial for pattern separation (Yamamoto et al., 2021).

Another line of research that focuses on cognitive aging also associates EC2a to pattern separation. We found that loss of EC2a recurrent inhibition in cognitive aging could account for reduced pattern separation, potentially through distorted TSP representations and altered activity levels. Consistent with this finding, it has been reported that diminished grid-cell-like representation (which would be a predicted consequence of low EC2a recurrent inhibition) in older adults is associated with poorer navigational ability (Stangl et al., 2018). In our model, age-related disinhibition in EC2a naturally leads to hyperexcitability in CA3, which lead to both reduced pattern separation and pattern completion in the face of incomplete cues. However, further empirical work would be needed to attribute underlying changes in EC2a neuronal circuits as direct causes of memory deficits in aging, because many other changes in the entire EC-HPC pathway could potentially be responsible (Burke & Barnes, 2006). Furthermore, additional work is necessary to more directly test the hypothesis that the previously-described pattern completion *improvements* in the aging population (Yassa & Stark, 2011) might instead be due to a response bias (Huh et al., 2006; Vieweg et al., 2015). A more strict definition of pattern completion, which we favor, applies only to “recovery of a stored pattern from a fragment” (Willshaw et al., 2015).

Studies have shown that LEC2a is among the earliest brain regions affected by Alzheimer’s disease (AD) pathology (Braak & Braak, 1991; Reagh & Yassa, 2017; Stranahan & Mattson, 2010), with a significant loss of Reelin+ neurons associated with the progression of AD (Gómez-Isla et al., 1996). More recent research suggests that increased intracellular expression of amyloid is selectively associated with Reelin+ neurons in LEC2a during the early stages of AD (Kobro-Flatmoen et al., 2016, 2021), pointing to the intertwined relationship between EC2a neuronal dynamics and AD. We simulated the loss of EC2a neurons and found that it roughly paralleled the progression of memory deterioration in AD, although multiple other factors could also contribute to this progress (e.g., loss of synapses from EC2a to DG). Finally, it is worth pondering whether reduced recurrent inhibition seen in the preclinical stage could also lead to progression of AD, as reduced grid-cell-like representation in young adults is related to genetic risk for AD, and excitatory and inhibitory imbalance may drive the pathogenesis of AD (Bi et al., 2020).

The RIDE model suggests several empirically-testable predictions. First, we predict that higher recurrent inhibition in Reelin+ neurons in EC2a would lead to better pattern separation in animal models, which could be tested behaviorally and neurally. Given that MEC and LEC are hypothesized to process different streams of cortical inputs while sharing similar underlying microcircuits (Witter et al., 2017), it is possible that weaker LEC2a recurrent inhibition could lead to object-based discriminability (Vandrey et al., 2020), which could be related to genetic risks for AD at an early stage (Kunz et al., 2015). Second, we predict that the TSP and the MSP serve different functions in not only episodic memory but spatial navigation, and information processing starts as early as in EC2a and EC2b/3, respectively. Specifically, the TSP supports pattern separation and pattern completion (Marr, 1971; O’Reilly & McClelland, 1994; Yassa, Mattfeld, et al., 2011), while the MSP facilitates the transformation of TSP representations into more structured cortical representations, promoting efficient generalization (McClelland et al., 1995; Schapiro et al., 2017). This could be tested by examining EC2a vs. EC2b/3 pattern similarities between similar environments.

The RIDE model also has several limitations. First, it does not account for the long-axis of the EC, which maintains a modular structure, particularly in MEC, with grid cell modules distributed along the long-axis (Hafting et al., 2005). Although one study has found that fast-spiking parvalbumin basket cells connect to more principal cells in dorsal vs. ventral MEC (Grosser et al., 2021), the biological basis of different modules remains to be characterized. Second, we only simulated EC2a neurons within a Continuous Attractor Network connectivity pattern, which has been used for simulating classical grid-cell coding, but there are other types of cells in EC2a that might not share the same underlying computational mechanisms with the ones we simulated. Third, we did not consider the “big-loop”, which connects the TSP with the MSP through connections between different layers of the EC. In the brain, these functionally separated pathways are likely to be more dynamically interacting and interdependent than is captured in this simplified model. Lastly, our model attributes the decline in lure discrimination among older adults to the loss of EC2a recurrent inhibition, but other changes in the TSP could potentially cause similar effects (e.g., loss of perforant pathway synapses (Burke & Barnes, 2006), reduced inhibition in DG and CA3 (Reagh & Yassa, 2017), etc.).

In summary, the RIDE model presents a biologically-based explanation for how recurrent inhibitory dynamics within EC2a support pattern separation. We argue against a systematically organized map for the EC2a neuronal population and propose that recurrent inhibition generates dissimilar representations for highly similar inputs in the EC2a. Finally, we suggest that neuronal dynamics in EC2a could be a key factor in understanding memory deficits in the aging and AD population.

## Methods

### Model architecture

The RIDE (Recurrent Inhibitory Dynamics in the Entorhinal cortex) model is primarily an extension of our previously developed HPC model, Theremin (Zheng et al., 2022). It inherits the CA1 error-driven learning algorithm from the ThetaPhase model (Ketz et al., 2013) as well as the CA3 error-driven learning algorithm from the Theremin model (Zheng et al., 2022) to achieve high-level pattern separation with effective learning. In RIDE, the original superficial EC layer (ECin) is separated into EC2a and EC2b/3 layers to better capture EC’s anatomy. However, the EC layers (2, 3, and 5) in RIDE is still a general EC that does not differ across the long-axis and the medial-lateral axis. The model is implemented using Emergent in Go and simulations are run on a server.

### Recurrent inhibition in EC2a

We implement recurrent inhibition between EC2a Reelin+ neurons following Couey et al. (2013). Specifically, each EC2a neuron connects with nearby EC2a neurons with a weight as follows:

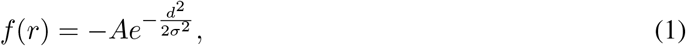

where *A* is the recurrent inhibition strength that ranges from 0 to 1, *d* is the distance between a receiving unit and a sending unit in the network, and *σ* is the standard deviation of the gaussian function. In the standard RIDE model, *max*(*d*) = 2 and *σ* = 4. Note that the gaussian weights also wraps, such that if the sending unit is at the margin of the layer, half of the receiving neurons will be on the other side of the layer with wrapping weights.

Unlike the traditional Continuous Attractor Network models (Burak & Fiete, 2009), RIDE does not implement a Mexican-hat connectivity profile. In a Mexican-hat connectivity pattern, neurons in the network exhibit excitatory connections with their nearby neighbors and inhibitory connections with more distant neighbors. This connectivity pattern resembles the shape of a sombrero (i.e., “Mexican hat”) with a peak of excitation surrounded by a ring of inhibition. The reason behind this implementation comes from neurophysiological evidence that there are few (Couey et al., 2013; Winterer et al., 2017) if any (Fuchs et al., 2016) excitatory connections between EC2a Reelin+ neurons. These neurons form strong disynaptic inhibitory connections with other Reelin+ neurons through Parvalbumin+ fast-spiking interneurons, giving rise to grid sheets similar to those formed by Mexican-hat networks (Couey et al., 2013). Note that this pattern exists in both MEC and LEC (Couey et al., 2013; Nilssen et al., 2018; Witter et al., 2017).

### Modeling the mnemonic discrimination task

In this paper, RIDE received a mnemonic discrimination task similar to the procedure in Reagh et al. (2018), where participants learned object-location associations on a computer screen and were later asked whether they had learned the object (object trial) or whether they had learned the object at a specific quadrant of the screen (spatial trial). To replicate the mnemonic discrimination task used in Reagh et al. (2018), our study created a set of stimuli that included 176 distinct patterns for target objects, each of which had a similarity of less than 50% with respect to the others. Additionally, we created 16 different patterns for spatial source quadrants, with each quadrant pattern having a similarity of less than 50% with respect to the others. As the original experiment had 42 trials and 4 quadrants, we aimed to ensure that each source pattern was presented equally, resulting in a total of 44 trials (i.e., 11 presentations per quadrant). To increase the difficulty of the task, we multiplied the number of patterns by four in our model.

The Input layer in RIDE has a 6-pool structure as in Theremin (Zheng et al., 2022), with the bottom 4 pools representing objects and the top 2 pools representing quadrants. Because the current paper focuses on object trials of the task, during the training phase the model were trained to auto-encode all the target object patterns randomly paired with spatial patterns. During testing, these same training patterns were shown again to test the model with target reconstruction. Additionally, object pools were flipped (i.e., turning on units that were off in targets, turning off units that were on in targets) systematically from 0% (identical) to 100% (fully dissimilar) to test the model with lure discrimination.

Although RIDE did not implement any mechanisms to directly obtain a final behavioral response as in the experiment, it measured two indices that are similar to the concept of hit rates (correctly identifying targets as “Old”) and correct rejection rates (correctly identifying lures as “New”). To obtain the proxies for these measures, we computed Target Reconstruction and Lure-Target Mismatch based on the degree of match (percent of EC5 neurons that matched the target patterns) between reconstruction patterns in the EC5 layer and input patterns given in the Input layer.

### Modeling the pattern completion task

In simulation 3, RIDE received a pattern completion task similar to the procedure in Vieweg et al. (2015), in which participants first learned associations between semantic labels and images (e.g., “dining room”) and were later tested on retrieving the corresponding label or responding with “new” for images with different levels of completeness (Fig 4A). RIDE was initially trained to autoencode binary random patterns representing different images. Later, it was tested on completing the missing bits from partial cues (which had increasing numbers of missing bits from the original training patterns) (Target) and completely new patterns (Foil).

## Acknowledgments

We extend our gratitude to Dr. Menno P. Witter for his insightful feedback and discussions on the anatomy of the entorhinal cortex. We also thank Dr. Rishidev Chaudhuri for his valuable discussions on the computational principles of the entorhinal cortex and grid cells.

## Funding Sources

RCO, CR were funded by the Office of Naval Research, grant numbers: N00014-20-1-2578. RCO was funded by the Office of Naval Research, grant numbers: N00014-19-1-2684, N00014-18-C-2067. The funders had no role in study design, data collection and analysis, decision to publish, or preparation of the manuscript.

## Competing Interests

R. C. O’Reilly is Director of Science at Obelisk Lab in the Astera Institute, and Chief Scientist at eCortex, Inc., which may derive indirect benefit from the work presented here.

## Footnotes

0 Although distributions of different cell types in MEC and LEC are more complicated than what we categorize here as EC2a and EC2b/3 (Ohara et al., 2019), we use EC2a to refer to Reelin+ cells that project to the HPC through the TSP, and EC2b/3 to refer to Calbindin+ cells in EC2b and Glutamate+ cells in EC3b that project to the HPC through the MSP (Nilssen et al., 2019).

1 Pattern completion is missing in the current simulation due to the single-epoch training regime we implemented in alignment with the experiment paradigm. Multi-epoch training that causes more long-term potentiation could be one way to bias the model, specifically CA3 with associative connections, towards pattern completion (O’Reilly & McClelland, 1994).

